# Anti-inflammatory immunomodulation for the treatment of Congenital Diaphragmatic Hernia

**DOI:** 10.1101/2023.11.07.565809

**Authors:** Mayte Vallejo-Cremades, Javier Merino, Rita Carmona, Laura Córdoba, Beatriz Salvador, Leopoldo Martínez, Juan. Antonio Tovar, Miguel Ángel Llamas, Ramon Muñoz-Chápuli, Manuel Fresno

## Abstract

Congenital diaphragmatic hernia (CDH) is a rare disease where the diaphragm does not develop properly altering lung development with no established therapy. We have analyzed the effect of anti-inflammatory immunomodulators that influence macrophage activation in animal CDH models. In the widely-used nitrofen-induced model of CDH in pregnant rats, administration of a single dose of atypical Toll-like Receptors TLR2/4 dual ligands (CS1 and CS2), 3 days after nitrofen, cured diaphragmatic hernia in 73 % of the fetuses, repaired the lesion with complete diaphragm closure. Moreover, they also improve pulmonary hypoplasia and vessel hypertrophy, enhancing pulmonary maturity of fetuses. CS1 treatment also rescued the CDH phenotype in the G2-GATA4^Cre^;Wt1^fl/fl^ CDH genetic mice model. Only 1 out 11 mutant embryos showed CDH after CS1 administration, whereas CDH prevalence was 70% in untreated mutant embryos. Mechanistically, CS1 stimulated the infiltration of repairing M2 macrophages (CD206+ and Arg1+) into the damaged diaphragm and reduced T cell infiltration. Alteration in retinoic acid pathways a have been also implicated in the etiology of CDH. TLR2/4 dual ligands also induced retinol pathway genes, including RBP1, RALDH2, RARα and RARβ, in the affected lungs and the diaphragm and in macrophages *in vitro*. The present results place atypical TLR2/4 ligands as a promising solution for CDH, where the own immune system of the fetus is responsible for repairing the hernia/damage in the diaphragm, ensuring the correct positioning and development of all organs.

## Introduction

Congenital Diaphragmatic Hernia (CDH) is a rare disease involving a defect in the diaphragm that frequently includes pulmonary hypoplasia. It occurs in about 1/3,000-5,000 of newborns (http://www.orpha.net/consor/cgi-bin/Education_Home.php?lng=EN#REPORT_RARE_DISEASES), being the cause of death of about 50,000 neonates per year worldwide, not counting the number of miscarriages caused by this disease. CDH is a birth defect resulting in the hernia of the diaphragm. Malformation of the diaphragm allows the abdominal organs to push into the chest cavity, hindering proper lung development. The presence of the abdominal contents in the thorax together with congenital alterations of pulmonary and cardiovascular development generate a picture of respiratory distress in the neonatal period due to variable degrees of hypoplasia and pulmonary hypertension, that are responsible for most deaths in children with CDH (reviewed in ^1–4^.

Current treatments for CDH are limited to the surgical repair of the diaphragm and reorganization of the organs in their proper position, combined with palliative interventions and treatments including pulmonary vasodilators, inotropic agents, surfactants, antibiotics, extracorporeal membrane oxygenation (EMCO), etc ^5,6^. As a result, the morbidity of survivors of this disease is very high and the quality of life very low. Despite this, the etiology of CDH largely remains unclear and currently is thought to be multifactorial. Multiple genetic factors along with environmental exposures and nutritional deficiencies have been proposed as possible etiologies for CDH. Although 20-40% of cases may have a genetic cause, the remaining ones are idiopathic in origin. Several studies have presented evidence of an increased risk for the development of CDH due to prenatal exposure to several maternal factors, such as alcohol, smoking, low vitamin A intake, obesity and antimicrobial drugs ^1^.

A widely accepted hypothesis to explain CDH relies on alterations in retinol metabolism leading to deficiencies in retinoic acid (RA) signaling, an important regulator of many genes during embryonic development. Vitamin A is essential for embryonic development and animal models genetically deficient in RA signaling and human vitamin A-deficiency are both associated with CDH induction ^7^. Moreover, many genes implicated in retinol metabolism are associated with CDH, which is consistent with the CDH retinoid hypothesis ^8^. This includes genes encoding proteins responsible for the cellular uptake of retinol (as STRA6, LRP2), cytoplasmic binding proteins of retinol and RA (as RBPs, CRABPs), enzymes involved in cellular RA metabolism (as LRAT, ALDHs), and nuclear RA receptors (RARα and RARβ) ^9,10^. In this regard, mutations such as those in the retinol receptor STRA6 ^11^ or in RALDH2 ^12^ have been described in humans with CDH. On the other hand, RA signaling pathway is an important regulator of diaphragm embryogenesis and defects in this pathway, or its downstream targets, can contribute to the development of Bochdalek hernia in CDH ^13^.

There is also evidence of genetic causality in patients with CDH. Although there are cases of familial CDH, most genetic data point to a multifactorial inheritance. In fact, multiple genes have been identified in patients with CDH, with COUP-TFII/Nr2f2, Friend of GATA2 (FOG2/zfpm2), GATA4, WT1 and SLIT3 being widely implicated (revised in ^9,10^. Results obtained with genetically deficient or knockout (KO) animals and animal models of CDH point to these same genes found in patients with CDH, besides others, most of them implicated in the development and differentiation of the diaphragm and/or lung, but they have yet to be corroborated in humans ^9,10^. On the other hand, there are several animal models that reproduce the symptoms of CDH ^14^. These include treatment during gestation with the herbicide teratogen nitrofen ^15,16^, being its teratogenic activity ascribed to alteration in the regulation of RA synthesis ^17,18^.

Macrophages are more than just professional phagocytic cells or antigen presenting cells in innate immunity or essential for the orchestration of adaptive immune response. Their functions include their homeostatic capacity to repair tissue after injury ^19^. Macrophages can be activated in different ways leading to M1 (classical) or M2 (alternative) differentiation ^20^. M1 macrophages are considered pro-inflammatory while M2 macrophages are anti-inflammatory and involved in tissue repair. An essential role is increasingly being given to macrophages in tissue remodeling and development during ontogenesis, either directly or through interaction with progenitor cells or stem cells ^19,21^. Large numbers of macrophages are present in almost all developing organs ^22^. In relation to CDH it has been described many years ago that macrophages infiltrate the fetal diaphragm in rats. These macrophages appeared to be involved in the removal and remodeling of muscle fibers in a homeostatic manner ^23^. RA metabolism is a property of M2 macrophages or dendritic cells (DC) activated by Toll Like receptor (TLR)2 ligands ^24,25^. Moreover, a recent report has shown that blockade of macrophage inhibitory factor (MIF) results in reduced macrophage migration and ameliorates pulmonary hyperplasia in a model of CDH ^26^.

We hypothesized that macrophages contribute to homeostatic development of the diaphragm and lung, and as a result immunomodulators can activate the transcription of several genes of the retinoic pathway and inducing M2 type differentiation macrophages. Retinoic pathway activation and M2 macrophages would in turn help repair damage caused by teratogens/mutations associated with CDH development and would contribute to tissue remodeling and organ development. To demonstrate this hypothesis, we tested several anti-inflammatory immunomodulatory candidates (TLR2 or TLR2/4 dual ligands) *in vitro* and on CDH animal models and found that they have a very significant curative effect.

## Results

### Effect of TLR ligands in nitrofen-induced CDH

Initially, we tested several TLR ligands in a rat model of CDH induced by nitrofen. Nitrofen was administered to pregnant rats at embryonic stage E9.5 to induce CDH and inoculated with several TLR ligands 3 days later (E12.5) to allow organ alterations to take place. On day 21 ^st^ of gestation, the rats were sacrificed and the embryos were examined for the presence or absence of diaphragmatic hernia (CDH+ or CDH-, respectively). Pregnant rats exposed to the herbicide nitrofen had between 60-85% of fetuses (73% on average in Figure 1A) with CDH. Treatment of the pregnant mothers with a single dose of TLR2/1 or TLR2/6 ligand, 3 days after nitrofen administration reduced on average CDH+ fetuses from 73 % in PBS treated animals to 46% and 52% respectively. This represents a relative decrease (or cure) in CDH incidence caused by nitrofen of 37% and 29 % respectively.

**Figure 1.**
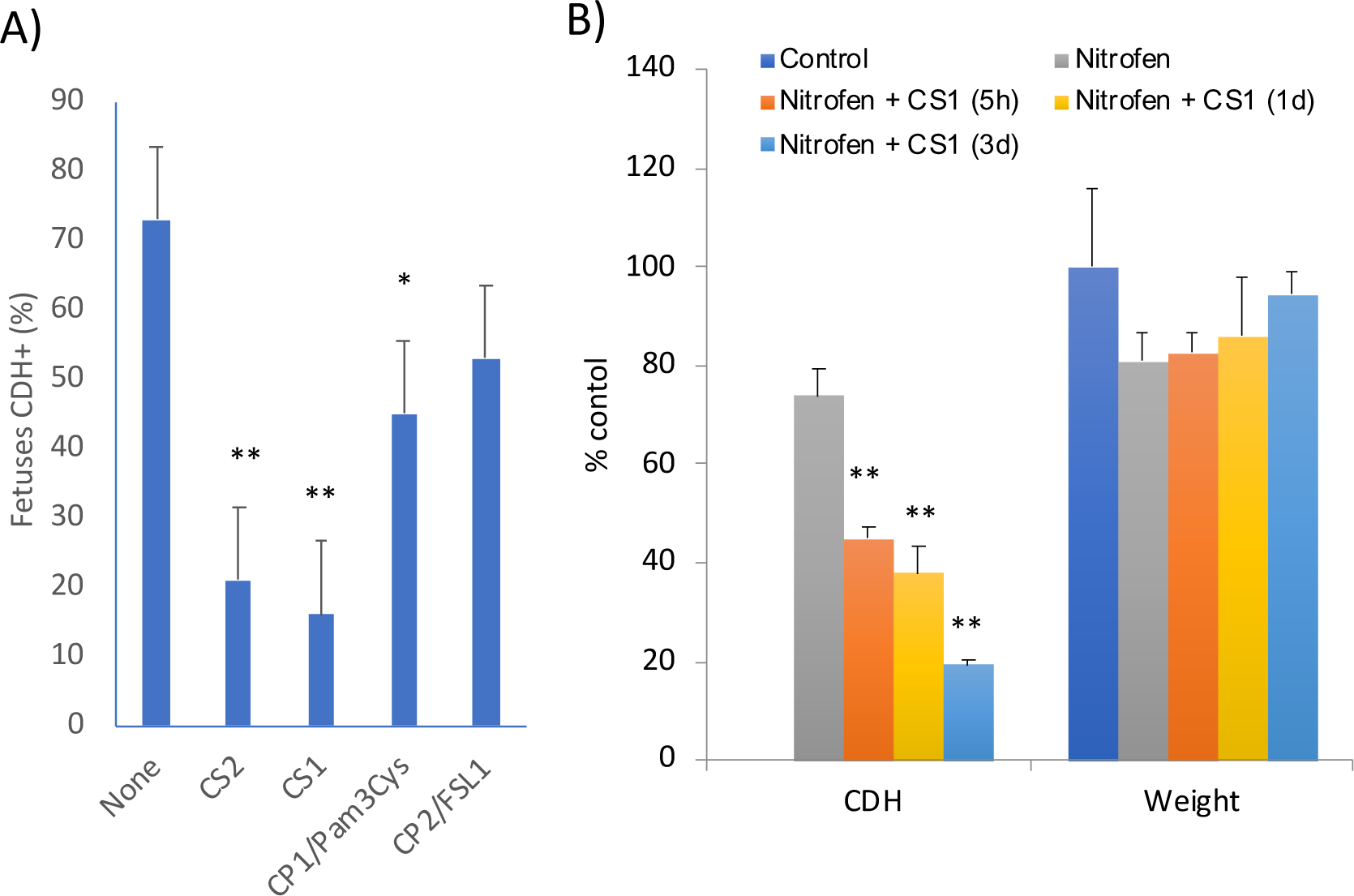
Effect of TLR ligands in diaphragmatic hernia induced by nitrofen. A) Pregnant Wistar rat females (4 per group) were administered nitrofen, or olive oil as placebo (control), at E9.5 and 3 days later injected intraperitoneally with 100 μg/Kg of the indicated immunomodulators or only saline. B) Pregnant Wistar rat females (3 per group) were administered nitrofen at E9.5 and 5h, 1 day or 3 days later injected intraperitoneally with 100 μg/Kg of CS1 or saline. Diaphragmatic hernia and weight of the foetuses were recorded at E21 as described in Methods. ** = p>0.001, * p>0.01 respect to untreated nitrofen group.

More importantly, atypical LPS which are a new class of dual TLR2-TLR4 ligands from soil bacteria (rhizobiales) ^27^ were much better at reducing this CDH incidence. LPS from *Rhizobium rhizogenes* K-84 (CS1) reduced CDH incidence to only 16% whereas LPS from *Ochrobactrum intermedium* (CS2) resulted in to 21% reduction This represents a relative decrease of 79% and 72 % in hernia generation respect to the nitrofen controls with saline treatment (Figure 1A). Treatment with LPS from *E.coli* was toxic to the fetuses, and no fetuses were recovered.

The histological analysis of fetus from pregnant mothers treated with CS1 at E18 and E21 (Figure 2) showed that in healthy fetuses, a complete diaphragm is clearly observed with a normal morphology and homogeneous muscle fibers separating the abdomen from the thorax (Figure 2A). As expected, fetuses from mothers that were administered nitrofen presented with an invasion of the thorax by the liver, due to partial or total absence of the diaphragm, combined with amorphous morphology. At E18 that the liver displaced the heart (C) and the left lung (LL), while at E21 the diaphragm was incomplete and showed histological alterations. Importantly, treatment with CS1 immunomodulator prevents damage to the diaphragm. For example, at E18 the pleuroperitoneal fold was incompletely covered by the muscle fibers (separating the abdomen from the thorax). At E21, most of fetuses of nitrofen administered mothers and treated with the immunomodulator, presented a complete diaphragm although the morphology was more disorganized and hypertrophied. Identical results were found with animals treated with CS2 (not shown).

**Figure 2.**
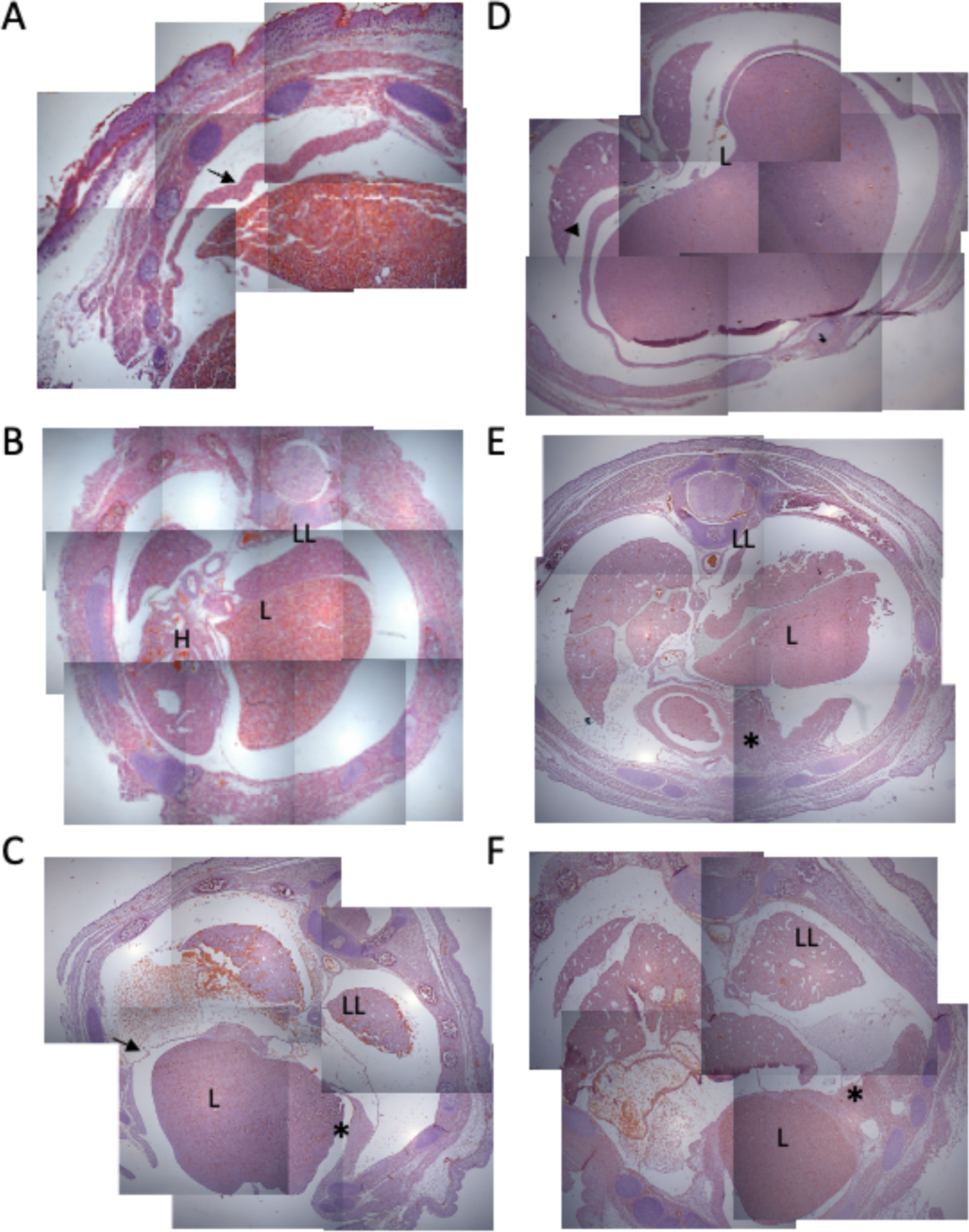
Representative images of rat foetuses of nitrofen-treated mothers with CS1 immunomodulators. Pregnant Wistar rat females were administered nitrofen, or olive oil (control), at E9.5 and 3 days later injected intraperitoneally with 100 mg/Kg of CS1 immunomodulator or only saline. Histological analysis of the foetuses were performed at E18 (A, B and C) and at E21 (D, E and F) as described in methods. A and D) healthy controls; B and E) nitrofen CDH+ (with hernia); C and F) nitrofen + CS1 CDH-, treated without hernia. In healthy foetuses diaphragm is clearly observed as a complete diaphragm and with a morphology of normal and homogeneous muscle fibers separating the abdomen from the thorax (arrow). In the nitrofen foetuses administered, the liver is invading the thorax, due to partial or total absence of the diaphragm that presents with amorphous morphology. At E18 (B) the image shows that the liver displaces the heart (H) and the left lung (LL), while at E21 (E) the diaphragm is incomplete and with alterations in its morphology (asterisks). Treatment with immunomodulator rescues the damage to the diaphragm. At E18 the pleuroperitoneal fold (arrow) is completely covered by the muscle fibres (asterisks) separating the abdomen from the thorax. The day 21 foetuses administered nitrofen and treated with the immunomodulator present a complete diaphragm, although the morphology is more disorganized and hypertrophied (asterisk).

We conducted trials with compounds CS1 and CS2 testing different dosages and times of administration. Again, we used the nitrofen model to produce CDH at day 9.5 of gestation and we treated with CS1, CS2 or saline at 5h, 1 or 3 days after nitrofen exposure. Hernia formation and weight of the embryos in each group were recorded. Interestingly, the ability of the compound CS1 (shown in Figure1B) and CS2 (not shown) in reducing CDH+ fetuses were increasingly higher from 5h to 3 days post-treatment. The curative effect was strikingly more efficient when the treatment with CS1 and CS2 immunomodulators was applied 1 day and especially 3 days after nitrofen administration, reaching then the maximum healing of the hernia. Embryo weight showed a slight increase following treatment with CS1 and CS2 TLR ligands compared to saline-treated rats, but this difference was not statistically significant (Figure 1B). Furthermore, no significant differences were detected when CS1 and CS2 were administered at different doses (data not shown). These findings suggest that as immunomodulators rather than classical drugs, CS1 and CS2 might influence the immune response largely in dose independent manner.

Alteration and retarded development of the lung is a characteristic of CDH ^1,3^. Histological analysis of the embryonic lungs showed clear differences among the groups treated with CS1 (Figure S1) and CS2 (Figure S2). Those from nitrofen CDH+ fetuses were notably smaller than the rest including the CS1-treated CDH+ fetuses (with herniation), and even smaller than CS1 or CS2-treated CDH-fetuses (without herniation). The average size of lungs in CDH+ nitrofen fetuses was approximately 23% smaller than lungs in fetuses treated with CS1. Furthermore, the lungs of CDH+ nitrofen fetuses showed altered morphology with less defined lobes. Importantly, CS1-treated fetuses exhibited lung morphology resembling that of normal mature lungs in 21-day-old fetuses with an increased number of respiratory bronchioles, alveolar ducts and alveoli (Figure S1A), while saline treated CDH+ fetuses displayed a more immature lung appearance, with underdeveloped and less clearly defined alveolar structures. Similar results were observed at E18 with the nitrofen-treated groups exhibiting more immature lungs compared to the control group (Figure S1B). The differences in alveolar ducts, alveoli, and respiratory bronchioles were evident as well. The untreated nitrofen group showed reduced airspace, especially in CDH+ fetuses, when compared to the treatment group and healthy controls. In contrast, the treatment group demonstrated better lung development in terms of size and an increased number of alveolar ducts especially in CDH-fetuses. Morphometric analyses were conducted to assess pulmonary vascular hypertrophy which contributes to pulmonary hypertension Healthy controls exhibited a highly stained media layer of the aorta wall with Van Giemson staining. In contrast, the nitrofen-CDH+ showed reduced staining of elastic fibers, indicating areas of tissue injury (Figure S1C-D). However, CS1 immunomodulatory treatment reversed this effect, supporting the beneficial impact of CS1 treatment in mitigating the negative effects of pulmonary hypoplasia and associated pulmonary vascular changes in CDH. Identical results were found with animals treated with CS2 (not shown).

### Effect of TLR ligands in WT1 conditional KO (G2-GATA4^Cre^;Wt1^fl/fl^) mice

Next, we decided to confirm the effects of CS1 treatment, using a genetic model, a mutant mouse with conditional ablation of the Wilms’ tumor suppressor gene (*Wt1*) in lateral plate cells expressing GATA4 under the control of the G2 enhancer ^28^. The use of this WT1 conditional knockout overcomes the early embryonic death caused by systemic deficiency of WT1 and it constitutes a valuable animal model of CDH. Defect in the post hepatic mesenchymal plate as well as a strong reduction of the septum transversum mesenchyme could be observed as early as E10.5 G2-Gata4^Cre^;Wt1^fl/fl^ mutant embryos. About 70-80% of these mutant embryos developed typical Bochdalek-type CDH, with diaphragmatic defect, liver invasion of the left pleural cavity and hypoplasia of the left lung ^28^.

We treated G2-GATA4^Cre^;Wt1^fl/fl^ pregnant mice mothers with CS1 or PBS intraperitoneally twice at E9.5 and E10.5. Embryos were analyzed at E15.5 (Figure 3 and S3). Very remarkably, CS1 treatment rescued the CDH phenotype in the G2-GATA4^Cre^;Wt1^fl/fl^ model. Of the 11 mutant fetuses analyzed, only 1 had diaphragmatic hernia (9% incidence) (Figure 3) when prevalence of the CDH+ phenotype in non-treated mutant embryos was 70% as described ^28^. In all but one CS1-treated mutants the pleural cavities were completely closed in contrast to the untreated mutant mice (Figure S3). Although the diaphragm does not appear normal, being thicker, more irregular and with less organized musculature, it is completely closed (Figure S4). The sinus venosus of all CS1-treated embryos was abnormal indicating that WT1 was deleted in all treated animals (Figure S3). This is an internal control to confirm the deletion of WT1 deletion and indicates that hernia was healed by CS1 and that the observed decrease on CDH was not due to a deficient deletion of the WT1 gene in those animals treated with CS1. It is also important to mention that control embryos treated with CS1 were normal (Figure 3).

**Figure 3.**
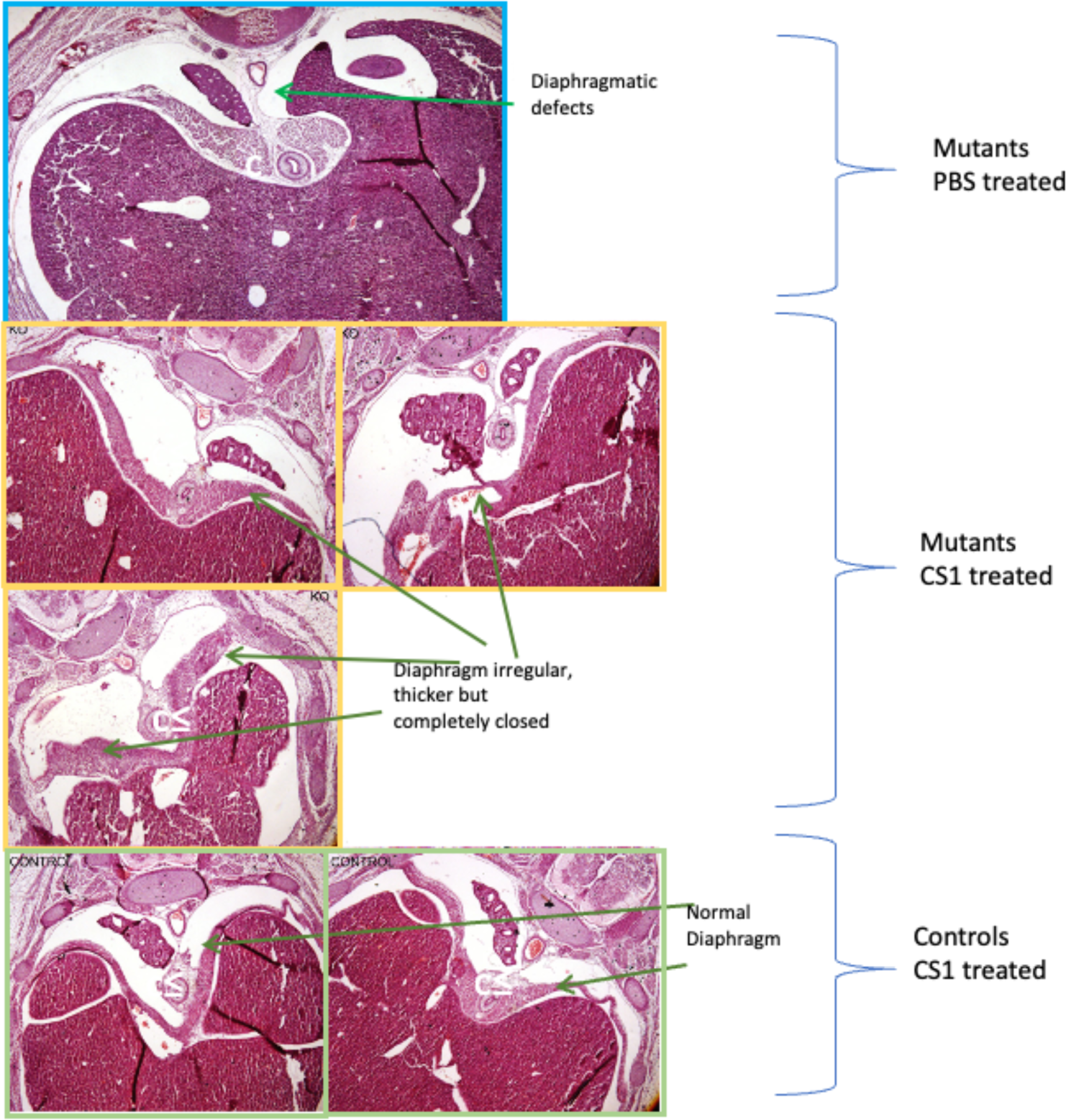
Effect of CS1 in G2-GATA4^Cre^;Wt1^fl/fl^ mice. Pregnant mice mothers were treated with 100 mg/Kg of CS1 or saline intraperitoneally, twice at E9.5 and E10.5. Embryos were analysed at E15.5. Images representatives from WT1 mutant embryo of mothers treated with PBS or CS1. Embryos from control not mutant mothers are also shown.

### Immune cell infiltration in damaged organs in CDH

The detection of immune cells is an essential step in order to understand the mechanism of action of the tested compounds and to determine what cells are activated through stimulation by immunomodulators such as CS1 and CS2. First, we measured expression markers of macrophages (CD68+), B (CD20+) and T (CD3+) lymphocytes and neutrophils (p67+) with immunohistochemistry techniques (Figure 4). In the CS1 treated groups, both CDH- and CDH+ fetuses were analyzed separately.

**Figure 4.**
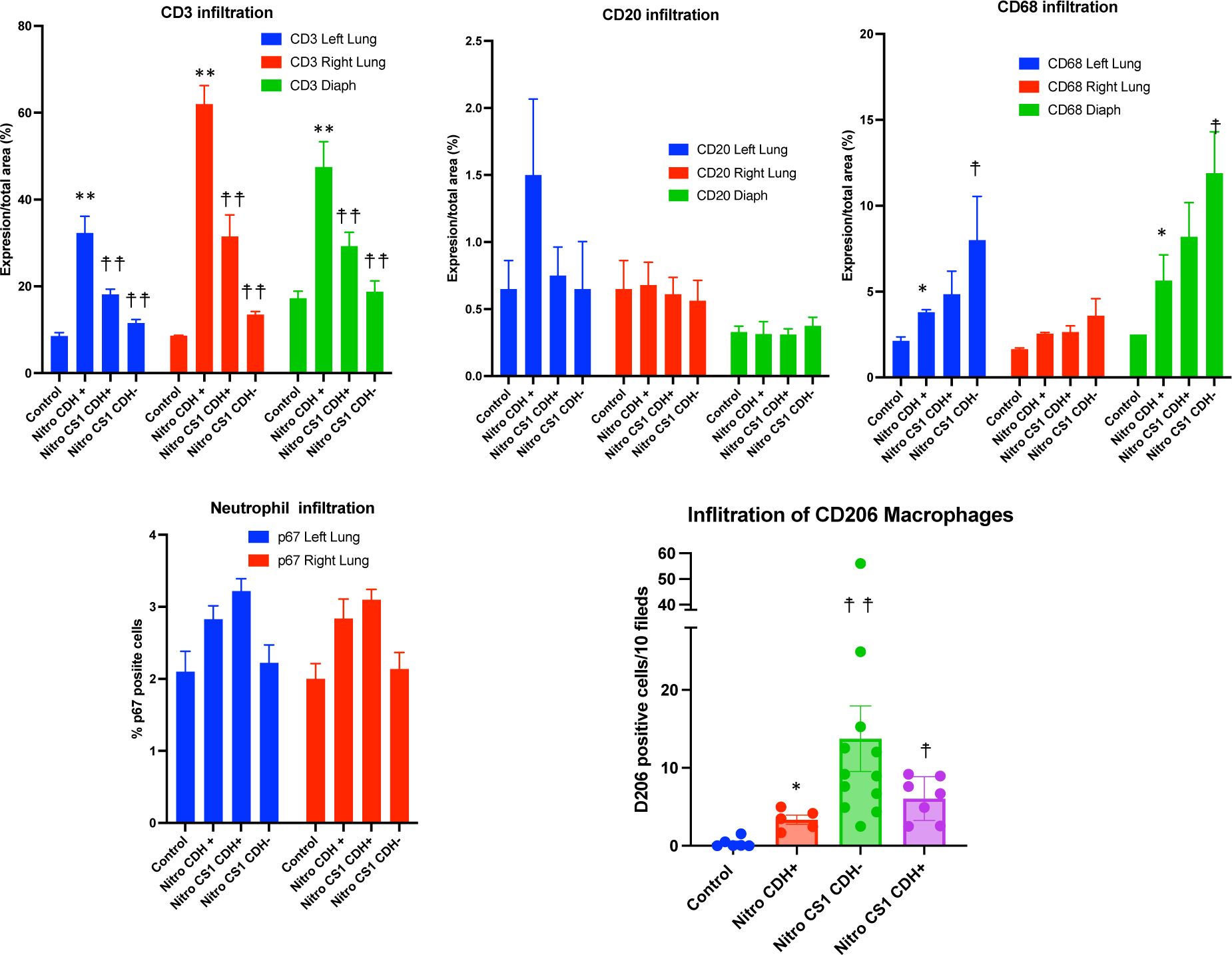
Immune infiltration in lung and diaphragm induced by nitrofen and CS1 treatments. Pregnant Wistar rat females (3 per group) were treated with nitrofen or olive oil (control) at E9.5 and 3 days later injected intraperitoneally or with 100 mg/Kg CS1 or saline. A-D) Indicated cell infiltration in the diaphragm, left and right lung was determined by immunohistochemistry at E21. In the CS1 treated groups, both CDH-negative and CDH-positive foetuses were analysed separately. E) CD206 expression was determined by immunofluorescence at E18. ** = p>0.001 and * = p>0.01, respect to control normal group; ☨ ☨ = p>0.001 and ☨ = p>0.01 respect to untreated CDH+ nitrofen group

In rats with CDH induced by nitrofen, a significant and high increase in CD3 T cell infiltration was observed in both the left and right lungs as well as in the diaphragm of the embryos at E21. However, in fetuses from pregnant rats treated with CS1, this infiltration was notably reduced, even in the few cases where the diaphragm did not close completely. This reduction was particularly remarkable in the cured CDH-fetuses (Figure 4). No relevant effect was observed in B cell infiltration (Figure 4). Examining the percentage of p67+ cells, a marker for neutrophils, in lung tissue, we found a lower number of p67+ cells in both the control and CS1-treated CDH+ groups (Figure 4). This indicates a higher neutrophil infiltration in CDH+ pups. No p67+ infiltrating cells were detected in the diaphragms.

Next, we examined the macrophage infiltration in the diaphragm. We observed a significant increase of CD68+ cells in all CDH+ groups compared to the control or CDH-groups. Interestingly, although nitrofen induced an infiltration of CD68+ macrophages in left lung and diaphragm, CS1 significantly increased this infiltration, especially in the CDH-fetuses (Figure 4). This suggests the recruitment of macrophages for tissue repair and resolution of damage in the two most affected organs, left lung and diaphragm as a result of CS1 administration. To corroborate this, we specifically studied the repairing macrophage population by using the M2 macrophage specific marker CD206 (Figure 4E). A great infiltration by M2 macrophages was observed in all nitrofen-treated animals. However, in CS1-treated CDH-animals the infiltration of this M2 population to the diaphragm was 2-3 fold higher than in nitrofen CDH+ animals. This indicates that there is an active recruitment of these M2 cells to facilitate the repair of the damaged tissue in CS1-treated CDH-animals.

### Mechanism of action of CS1 and CS2 in CDH

To gain a better understanding of the effects of CS1 in tissues, we analyzed the gene expression patterns in rats exposed to nitrofen and treated with CS1. RT-PCR was performed to analyze gene expression in the lungs, diaphragm, and spleen. Figure 5 provides a summary of two independent experiments, each including all fetuses from pregnant rats (three rats per group in each experiment) treated with nitrofen at E9.5, with or without administration of CS1 at E12.5 and analyzed at E21. The results were normalized for comparison with the untreated control group using real time QC-PCR. Fetuses from control animals treated with CS1 at E12.5 showed no significant changes in gene expression at E21 compared to fetuses from untreated control rats (not shown). Nitrofen administration alone induced minor changes in gene expression in the analyzed organs at E21. The most notable changes were the induction of *Arg1* in the spleen and diaphragm, *Ccl2* in the diaphragm, and *Wt1* and *Stra6* in the lungs. Interestingly, treatment with CS1 of nitrofen animals resulted in a significant increase in *Arg1*, *Aldh1a2*, *Rbp1* and *Rarb* in the diaphragm. The spleens from fetuses of CS1-treated rats also showed increases in *Arg1*, *Aldh1a2*, *Rara*, *Ccl2*, *Pparg*, and *Slit3*. In the lungs, however, only *Arg1* showed a significant and high induction following CS1 treatment, Additionally, there was a tendency for increased expression t in retinoic pathway genes *Aldh1a2, Rara* and *Rarb*, while the inductions of *Stra6* and *Wt1* caused by nitrofen were reduced. These results indicate a significant increase in Arg1 and the RA pathway, along with a decrease in *Wt1* and *Stra6* due to CS1 treatment in the affected organs and spleen.

**Figure 5.**
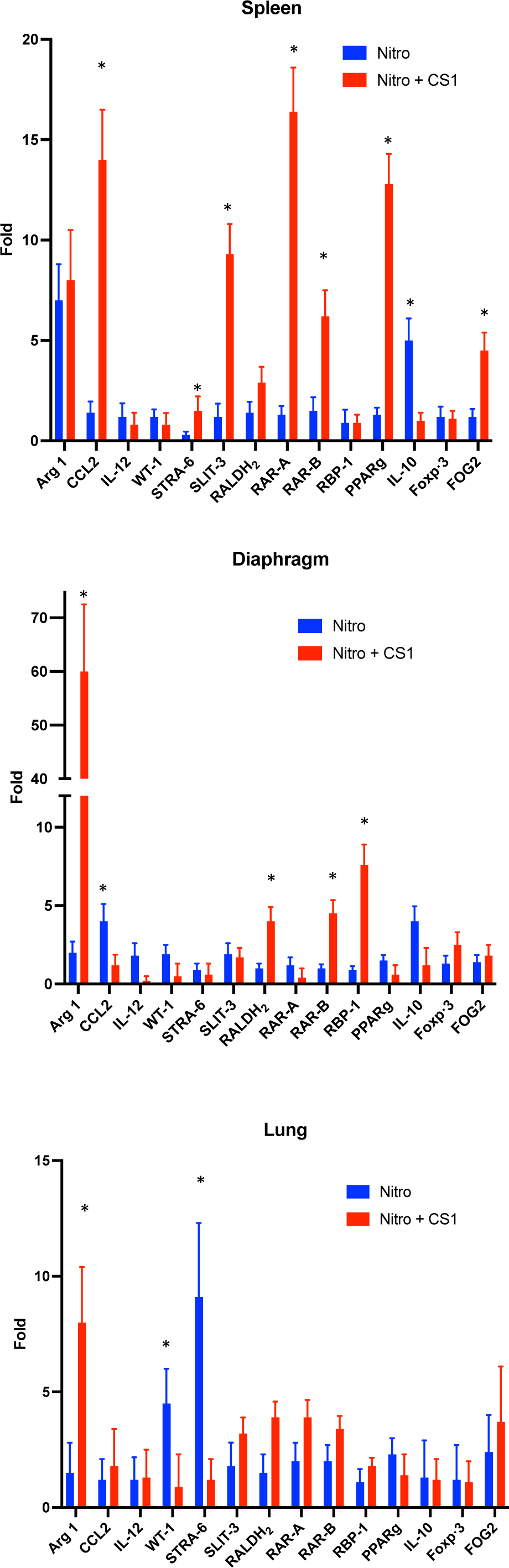
Effect of TLR ligand on gene expression in vivo. Pregnant Wistar rat females (3 per group) were treated with nitrofen or olive oil (placebo) t at E9.5 and 3 days later injected intraperitoneally with 100 mg/Kg CS1 or saline. RT-PCR of the indicated organs was carried out at E21. Results are the mean of 2 independent experiments with 5 foetuses per group each.

### CS1 and CS2 ligands induce retinoic pathway genes in macrophages

To further understand the effect of the treatments in the immune activation, we investigated the ability of CS1 and CS2 to activate macrophages (Figure 6). CS1 and CS2, induced *Aldh1a2* and *Rbp1*. Additionally, both induced *Arg1,* a marker of M2 macrophages and also weakly induced *Slit3* which has recently been found to be also expressed in M2 mouse peritoneal macrophages ^29^. Interestingly, by using macrophages from TLR2 or TLR4 deficient macrophages, we found that those effects were dependent on both TLR2 and TLR4 receptors. In addition to those genes of the retinol pathways, both CS1 and CS2 also induced *Ccl2* and *Il12*, being this effect dependent more on TLR4 than on TLR2. These *in vitro* effects align with the observed infiltration of macrophages and gene expression profile primarily observed in the diaphragm, but also in spleen and lungs, of the CS1-treated fetuses shown above.

**Figure 6.**
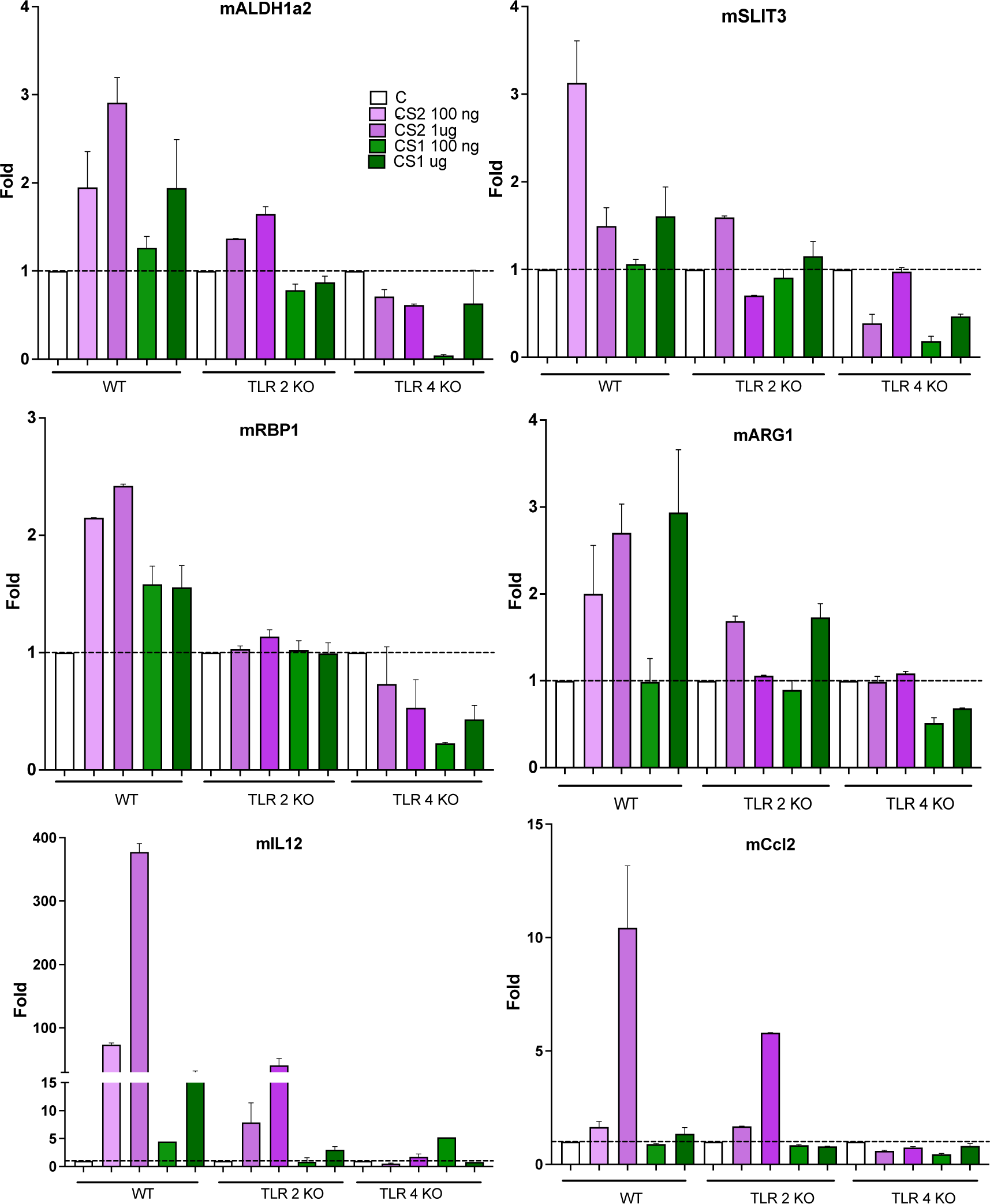
Effect of TLR ligands on gene expression by macrophages. Peritoneal macrophages from WT, TLR2 or TLR4 deficient mice were stimulated with CS1 and CS2 at the indicated doses and 14 hr later the gene expression was analysed by RT-PCR. Results are the mean of 3 independent experiments in duplicate.

## Discussion

In this study, we demonstrate the efficacy of a new class of anti-inflammatory immunomodulators, atypical LPS, for the treatment of CDH in both, teratogen-induced and genetic mouse models. Notably, we observed successful CDH rescue in a genetic model of CDH, the WT1-deficient mice with ablation of Wt1 in the territories giving rise to the diaphragm, making it the first instance, to our knowledge, of immune modulation rescuing an inborn genetic defect. Our findings show that the CS1 and CS2 ligands, which modulate TLR2/4, can effectively redirect the retinoic acid pathway and the innate immune system in CDH promoting tissue repair and remodeling. Specifically, these TLR dual ligands induce the infiltration of M2 macrophages, particularly in the diaphragm, and significantly reduce the number of hernias in the fetuses of pregnant females after CDH induction with nitrofen. Moreover, CS1 and CS2 treatment not only reduces the incidence of hernias in both CDH animal models but also improves pulmonary hypoplasia and vessel hypertrophy, enhancing pulmonary maturity of fetuses. Additionally, CS1 treatment induces the expression of retinoic acid pathway genes in damaged organs. Overall, our study highlights the promising therapeutic potential of CS1 and CS2 ligands as immune modulators for the treatment of CDH.

Toll-like receptors (TLRs) recognize endogenous stress signals and play an important role in many processes but especially in myeloid biology ^30^. Macrophages can be activated in response to different TLR ligands and undergo M1 (classical) or M2 (alternative) activation ^20^. M1 is implicated in inflammation and pathogen elimination, and M2 in reparation, tissue remodeling, and wound healing. Arginase I (Arg1) is predominantly expressed in myeloid cells being a marker characteristic of M2 macrophages as it is the expression of CD206. Interestingly, in CDH fetuses we found higher levels of Arg1 in damaged organs and greater infiltration of CD206+ cells in the diaphragm after CS1 treatment suggesting that CS1 induces infiltration of M2 macrophages in the damaged organs.

Treatment with TLR2/4 ligands could result in a dual action beneficial for CDH treatment. Firstly, these ligands activate repairing macrophages favoring their migration to the damaged organs where they will decrease tissue damage while favoring remodeling, therefore allowing a better formation of tissues even if they were damaged. This concept aligns with studies have showing that blocking MIF, a cytokine that inhibits macrophage migration and therefore its blockade would favor macrophage infiltration, can improve pulmonary hyperplasia in CDH models ^26^. Secondly, immunomodulators promote *in situ* synthesis of RA pathway molecules such as RALDH2 which will increase RA levels favoring correct organ development. Combining these two actions, promoting repairing macrophages and increasing RA levels, could potentially enhance the overall therapeutic effect for CDH treatment.

Although not well recognized, the role of macrophages in development and ontogenesis is known. Macrophages are more than just professional phagocytic cells or antigen presenting cells essential in the orchestration of adaptive immune responses. An essential role is increasingly being given to the homeostatic capacity of macrophages in tissue remodeling during ontogenesis ^19^. Besides, they interact with progenitor cells or stem cells and this interaction may contribute to tissue repair and remodeling during ontogenesis ^19,21^. Remodeling and repair occur dynamically during ontogenesis and inflammation, and these processes are orchestrated by macrophages. Large numbers of macrophages are present in almost all developing organs ^22^.

An important function of macrophages in ontogenetic development is their trophic role since they produce a wide range of key trophic factors for the development of all tissues ^19^. They also contribute in a very important way to vascular development^31^. All of these factors have been implicated in one way or another in experimental models or patients with CDH. Macrophages are also required to eliminate apoptotic cells produced during normal development of various tissues ^32^. It has been described many years ago that macrophages infiltrate the fetal diaphragm in rats ^23^. These macrophages are frequently observed in the diaphragm during gestation (from day 16) and up to 2-4 weeks of age and then disappear and appeared to be involved in the removal and remodeling of muscle fibers in a homeostatic manner.

Interestingly, transcriptional analysis of macrophages isolated from mouse embryos and human fetal liver revealed that they had an M2 pattern consistent with the trophic function and tissue remodeling that has long been attributed to mononuclear phagocytes during development ^33,34^. Fetal M2 macrophages even exhibit an enhanced remodeling and wound healing phenotype relative to adult mouse M2 ^35^. In addition, increased expression of M2 markers (Arg1, Ccl17 and CD206) has been described in mouse lung development, which correlates with the stage of alveolar development, alveolar wall remodeling and microvascular maturation ^36^. Of those M2 markers, Arg1 and CD206 are augmented by CS1 treatment.

TLR2 and TLR4 differentially modulate M1/M2 differentiation, with TLR2 ligands capable of differentiating towards M2 macrophages and TLR4 towards M1 ^37,38^. Recently, we described that atypical dual TLR2/4 ligands will skew macrophages to M2 differentiation, without being proinflammatory ^27^. Moreover, TLR2 activation induces the enzymes that synthesize vitamin A, mainly RALDH2, and promotes signaling by RAR in myeloid cells ^25^. Likewise, RAR modifies the signal produced by TLR2 towards a monocyte capable of remodeling the extracellular matrix ^39^ and with M2 properties ^40^.

We observed an increase in several genes involved in the RA pathway upon treatment with atypical TLR2/4 ligands, CS1 and CS2. This effect was evident both in macrophages *in vitro* and, notably, in the organs of nitrofen-affected embryos *in vivo*. Specifically, we found an upregulation of *Aldh1a2* that encodes RALDH2, a key enzyme in RA pathway which is expressed in developing lungs and catalyzes the synthesis of RA. Interestingly, previous research has shown that nitrofen inhibits RALDH2 activity in CDH induced models ^41^. Additionally, we observed an upregulation of RBP1 by CS1 and CS2, the carrier protein involved in the transport of retinol in the cytoplasm before its conversion into RA by RALDH2. Both effects will lead to an increase in RA secretion by macrophages, similar to the induced *in vitro* by a TLR2 ligand ^25^. Those CS1 and CS2 activities were dependent of both TLR2 and TLR4, consistent with their described abilities to induce cytokine secretion of those types of atypical LPSs ^27^.

RA receptor genes (*Rara* and *Rarb*) have been linked to CDH ^41,42^. Knockout mice lacking these genes show significant lung hypoplasia and diaphragmatic defects ^43^. Interestingly, these genes are upregulated in CS1 treated groups, suggesting CS1 treatment may contribute to the restoration of transcription of important genes in the RA pathways with a central role in the CDH recovery process.

Vitamin A plays an essential role in the embryonic development and its deficit may lead to CDH as shown in pregnant rats fed with a vitamin A-deficient diet ^42^. It has further been reported that knockout of RBP (retinol binding protein) in mice significantly reduces retinol levels and causes embryonic abnormalities including pulmonary hypoplasia ^44^. Moreover, retinol and RBP levels are decreased in human newborns with CDH even though maternal RBP levels were comparable between mothers of CDH patients and mothers of healthy children ^45^. This suggests a specific defect in retinol homeostasis in the child rather than a maternal deficiency. Moreover, although nitrofen exposure causes a high incidence of CDH, coadministration with RA has a protective effect ^46^. Therefore, the observed upregulation of some RA pathway genes in CS1 and CS2 treated groups suggests that these ligands have the potential to increase RA secretion in the lungs, potentially contributing to the therapeutic benefits of CS1 and CS2 treatment in ameliorating CDH.

A recent study has shown that in mice TLRs are expressed in macrophages as early as E10.5 and that TLR ligands induce in these macrophages the ability to phagocytose apoptotic cells and secrete a broad spectrum of cytokines and chemokines ^47^. Thus, embryonic macrophages can be receptive to TLR ligands and this may explain some of the effects we observed after treatment with CS1 and CS2.

An essential consideration is that the CS1 and CS2 treatment is effective up to at least three days (at E12,5) after the induction of the teratogenic damage in rats or after the induction of the WT1 loss in the conditional WT1 mutant model. This finding is of great clinical significance because this embryonic stage (E12.5) in rats corresponds to over 20 weeks of gestation in human fetuses ^48^ and it is typically around this human pregnancy stage when the diaphragmatic hernia is first detected in a routine echography ^5,6^. Therefore, the immunomodulatory treatment described here, could potentially be used in human CDH cases, as effective treatment may be initiated at the time of diagnosis. Therefore, the favorable treatment window observed in the rat model provides promising prospects for clinical applications in human CDH treatment.

CDH lacks a etiological treatment, and current clinical management combined pharmacological and surgical measures, aiming to reduce the pulmonary hypertension and to close the diaphragm repositioning the affected organs into the abdomen; however, the overall survival remains alarmingly low ^6^. Unfortunately, the complex and multifactorial nature of the disease, with no single genetic cause, makes genetic therapies basically impractical. In this context, targeting the immune system to promote tissue repair in CDH represents an innovative and promising avenue of research. The findings of this study, demonstrating the effectiveness of CS1 and CS2 immunomodulators in rescuing CDH in both teratogen-induced and genetic mouse models, present a significant breakthrough in the field. The potential application of these immunomodulators in human CDH treatment, particularly during the early stages of gestation, holds great promise for improving outcomes and addressing the pressing medical need in this challenging condition. In support of this, the TLR2/4 dual ligand CS1 has been recently awarded the orphan drug designation by the European Medicines Agency (EMA).

## Material and Methods

### TLR Ligands

LPS from *Rhizobium rhizogenes* K-84 (CS1) and *Ochrobactrum intermedium* (CS2) were purified and characterized as previously described ^27^. The purity of those compounds was assessed by mass spectrometry with a purity level higher than 98%. LPS from *E. coli* O111:B4 was from Sigma. TLR2/TLR1 ligand Pam3CSK4 and TLR2/6 ligand FSL-1 were from InvivoGen: All LPS were resuspended in sterile PBS 1x.

### Nitrofen Rat model

Female Sprague-Dawley rats weighing 220-250 g and males with proven fertility, were housed in the facilities of the Research Unit of the Hospital Universitario La Paz in Madrid. They were provided with a special diet for rats and had access to water “*ad libitum*”. For controlled fertilization, females were caged with male (3×1) during one night after 24 hours of visual and olfactory contact. The time at which the vaginal smear demonstrates the presence of spermatozoa was considered day 0 of gestation ^18^.

The pregnant female rats were randomly divided into two experimental groups: the nitrofen group and the control group. On day 9.5 of gestation, the nitrofen group was administered 100mg of nitrofen (2,4-dichloro-4’-nitrodiphenyl ether, Sigma-Aldrich) in 1mL of olive oil via intragastric administration ^16^. In contrast, the control group received an equivalent volume of olive oil (placebo) using the same administration method. It is assumed that the percentage of fetuses with CDH after teratogen administration will reach more than 60% in each litter. Subsequently, on different days as indicated, the nitrofen group was further divided into two subgroups: one subgroup was intraperitoneally injected with the immunomodulators at a dose of 10μg/Kg, and the other subgroup received an equal volume of saline, serving as the non-treated group. On gestational day 18 or 21, the pregnant rats were euthanized by intracardiac injection of potassium chloride, and all the fetuses were carefully collected by caesarean section. Under a stereomicroscope, the fetuses’ diaphragms were examined for the presence of hernias. Based on the presence or absence of hernias and the treatment received, the fetuses were categorized into five groups for subsequent analysis: the control group (n=3), nitrofen with hernia group (Nitro CDH+, n=5), nitrofen without hernia group (Nitro CDH-, n=5), nitrofen with immunomodulator treatment (T) and hernia group (Nitro + T CDH+, n=5), and nitrofen with treatment but no hernia group (Nitro + T CDH, n=5). Some of the fetuses were stored in 10% buffered formaldehyde during seven days for future paraffin fixation and from the remaining lungs, spleens and diaphragms were separated and frozen at -80°C for RT-PCR.

Throughout the process, the animal experimentation followed ARRIVE guidelines and received approval from the Fundación para la Investigación Biomédica del Hospital Universitario La Paz ethics committee. The protocols were in accordance with Spanish legislation (RD 53/2013) and European legislation (2010/63/EU).

### Mouse lines and embryo extraction

The mice used in our research program were handled in compliance with the institutional and European Union guidelines for animal care and welfare. The procedures used in this study were approved by the Committee on the Ethics of Animal Experiments of the University of Malaga (procedure code 2015–0028). All embryos were staged from the time point of vaginal plug observation, which was designated as E0.5. Embryos were excised and washed in PBS before further processing.

The G2-GATA4^Cre^;Wt1^fl/fl^ mice model for CDH was previously described ^28^. In short, we used a driver based on the G2 enhancer of the Gata4 gene that drives expression of Gata4 in the lateral plate mesoderm from the stage E7.5, and by the stage E9.5 is active in the septum transversum and proepicardium, ceasing its activity by E12.5. The activity of this enhancer is completely absent in the intermediate mesoderm. In this model, it has been shown that WT1 is involved in the generation of the mesenchyme of the ST/PHMP/PPFs continuum through epithelial-mesenchymal transition. Those mice when crossed with the Wt1 floxed mice *WT1^fl/fl^* which allow the expression of the Cre recombinase and subsequent deletion of WT1 only on those tissues and in those days. Defect in the PHMP in G2-Gata4^Cre^;Wt1^fl/fl^ mutant embryos could be observed as early as E10.5. It provides a good model on the genesis of the Bochdalek hernia (70-80% of *G2-Gata4^Cre^; Wt1^fl/fl^* embryos developed typical Bochdalek-type CDH) ^28^. In those mice, strong reduction of the ST mesenchyme, in embryos was observed at E10.5. Liver invades left pleural cavity, and this lung becomes hypoplasic and diaphragmatic defects appeared lacking the continuity.

Pregnant females at E9.5 and E10.5 were administered a single intraperitoneal injection (IP) of CS1 diluted in PBS at a dose of 10 µg/kg of animal weight. Embryos were isolated at E15.5. Whole mount embryos were fixed in 4% paraformaldehyde in phosphate-buffered saline (PBS) at 4°C during 4-5 hours and processed for paraffin embedding. Hematoxylin-Eosin stain was performed using routine protocols.

### Isolation of Mouse Peritoneal Macrophages

C57BL/6 WT, TLR2 and TLR4 KO mice littermates were obtained from S. Akira. All mice were bred and maintained in the animal facilities of the Centro de Biología Molecular Severo Ochoa in Universidad Autónoma de Madrid. All animal procedures were performed in strict accordance with the European Commission legislation for the protection of animal used purposes (2010/63/EU). The protocol for the treatment of the animals was approved by the Comité de Ética de la Dirección General del Medio Ambiente de la Comunidad de Madrid, Spain (permits PROEX 128/15). Thioglycolate-elicited peritoneal macrophages (PM) were isolated from 6-8-week-old pathogen-free mice. Cells were cultured in RPMI 1640 (2 mM L-glutamine, antibiotics 100 units/mL penicillin, 100 mg/mL streptomycin) with 5% FBS and seeded into 6-well-pates at a density of 1×10^6^ cells/well. Cells were allowed to adhere for 2 h and then the medium was changed to remove non-adherent cells. After 24 h, medium was replaced with new complete medium prior treatment with TLR ligands ^27^.

### Morphometric and immunohistochemistry studies

After fixation, the fetuses were embedded in paraffin and 5 μm thick sections were cut with a microtome (Leica Biosystems). Morphometric analyzes were performed on E18 and E21 slides, stained with Hematoxylin/Eosin as described ^49^ from organs previously extracted and set in paraffin. The expression of immune system markers were mostly studied with inmunohistochemistry using the following antibodies: mouse anti rat CD68 (dilution 1:300 BIORAD, ref. MCA 341R) for monocytes/macrophages, rabbit anti CD3 (dilution, 1:400; Biorbvt, ref. orb10313) for lymphocytes T, rabbit anti CD20 (dilution, 1:400; Bioss, ref. bs-0080R) for lymphocytes B, rabbit anti mouse CD206 (dilution, 1:200; Bio Orbyt, orb180464) and rabbit anti human p67 (dilution 1:1000, Bioss antibodies). As secondary antibodies, Anti-Rabbit HRP (Millipore) for CD20 and CD3 and Anti-mouse HRP (Millipore) for CD68.

The samples were deparaffinized and pretreated to pH6 using PT-link (Dako-Agilent). Subsequently, they were incubated for 30 minutes hydrogen peroxide solution, to block the activity of endogenous peroxidase, and after that they were incubated for one hour with blocking solution (TBS+10% NGS+ 1% BSA + 0, 01% Triton X-100) to block non-specific binding of antibodies. They were then incubated with the corresponding primary antibody at 4°C overnight. Incubation with the secondary antibodies, was performed at room temperature for 45 minutes followed by three 5-minute washes in TBS. The signal was revealed with 3,3’-Diaminobenzidine (DAB) substrate (Palex) and the nuclei were counterstained with Mayer hematoxylin (Merck). The incubations were carried out in humid chambers to prevent the samples from drying out. Once the staining process is finished, the slides were dehydrated in an increasing alcohol gradient and finally the samples were mounted using DPX (Merck). Positive cells have a brown staining indicative of the precipitate of the DAB substrate. In order to separate non-specific from specific labeling in the immunohistochemical tests, negative controls were performed, in which the primary antibody was not included and therefore any labeling observed is considered non-specific.

The images were obtained using the Microscope Olympus BX41 and the software ImagePro plus 5.3 and analyzed with Image J program. For immunohistochemistry evaluation, three different areas of each lung and five different parts of the diaphragms were chosen for analysis in 5 fetus of each group and a “positive cell number/total cell number ratio” was stablishedElastica Van Giemson was used to study the morphometry of the vessels, particularly of the aorta artery in E18 thoracic sections in 5 fetus of each group. The width of the aortic adventitia was measured as a sign of hypertension. Increase of the wall thickness to vessel radius ratio (w/r) is related with increasing systemic pressure ^50^.

Confocal immunofluorescence was performed basically as described ^51^. Diaphragms, 3 fetuses per group, were fixed in 4% paraformaldehyde in PBS solution, incubated in 30% sucrose solution, embedded in Tissue-Tek O.C.T. compound (Sakura), and frozen in liquid nitrogen. Sections that were 10- to 15-µm thick were fixed in acetone. Incubation with the following antibodies was done at 4°C: 10 µg/mL goat anti-mouse CD206. Images were obtained using an LSM510 Meta confocal laser coupled to an Axiovert 200 (Zeiss). Four different parts of the diaphragms were chosen for analysis.

### mRNA Isolation and RT-qPCR

RT-qPCR was basically performed as described ^52^. Briefly, total RNA was extracted from each organ extracted, previously frozen at -70⁰C, using NZyol Reagent (NZYTech). cDNA was prepared by reverse transcription (GoTaq 2-Step RT-qPCR System, Promega) and amplified byPCR using SYBR® Green PCR Master Mix and ABI Prism7900HT sequence detection system (Applied Biosystems), with the primers shown in Table S1. The 2−DDCt method was applied to analyze the relative changes in expression profiling and all quantifications were normalized to the housekeeping gene RPL13A

For mouse peritoneal macrophages, cDNA was prepared by reverse transcription (GoTaq 2-Step RT-qPCR System, Promega) and amplified by PCR using SYBR® Green PCR Master Mix and ABI Prism 7900HT sequence detection system (Applied Biosystems). Primers used for qPCR analysis are listed in Table S2.

#### Statistical analysis

One way ANOVA was carried out, with a Bonferroni’s post-test for multiple Analysis was performed using GraphPad Prism 5 software. Quantitative results are expressed as means ± SEM or mean ±SD. Statistical analysis was performed using one-way ANOVA followed by Bonferroni multiple comparisons. A p-value less than 0.05 was considered statistically significant.

---

**Table S1.**
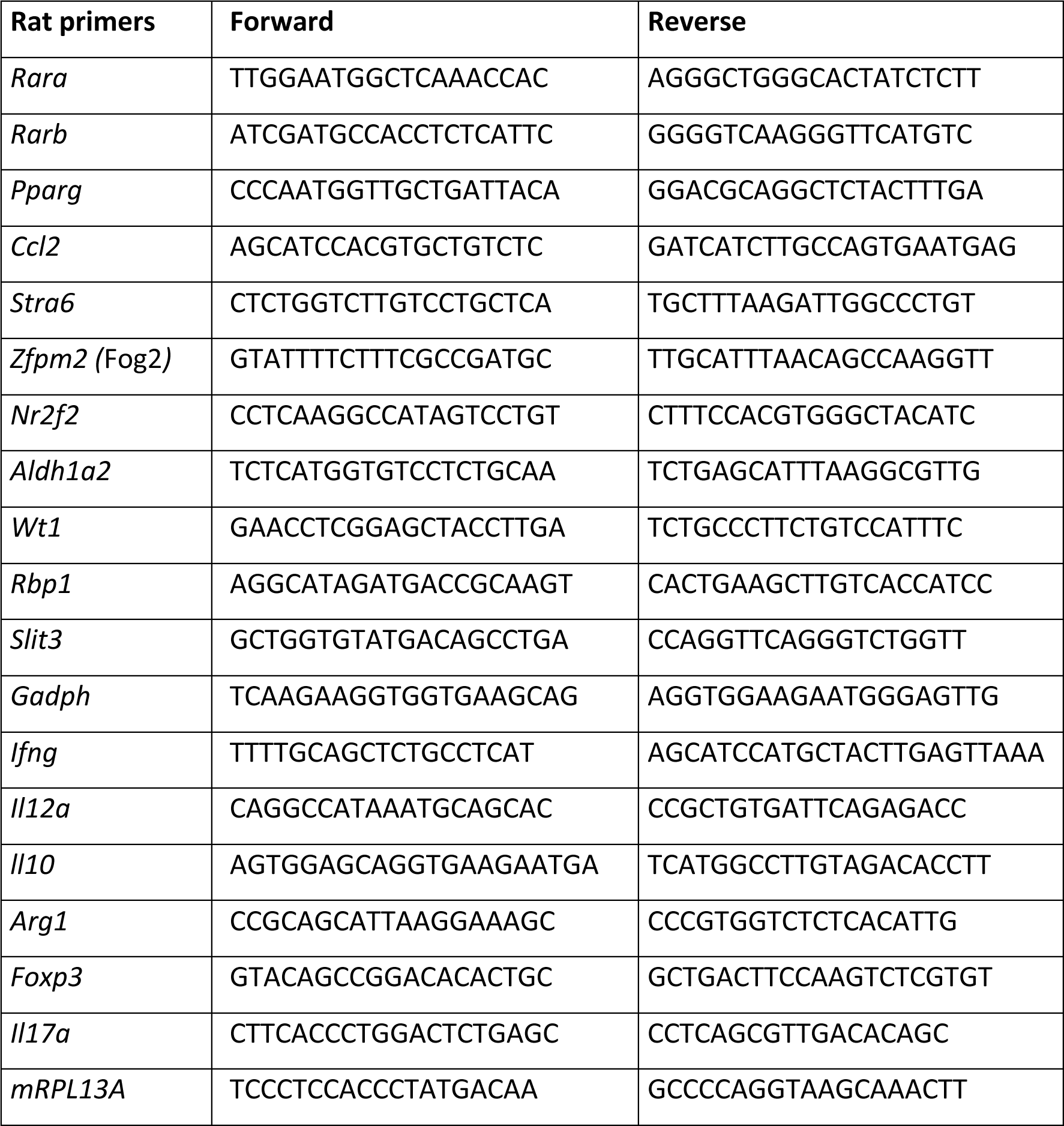
Rat RT-PCR primers.

**Table S2.**
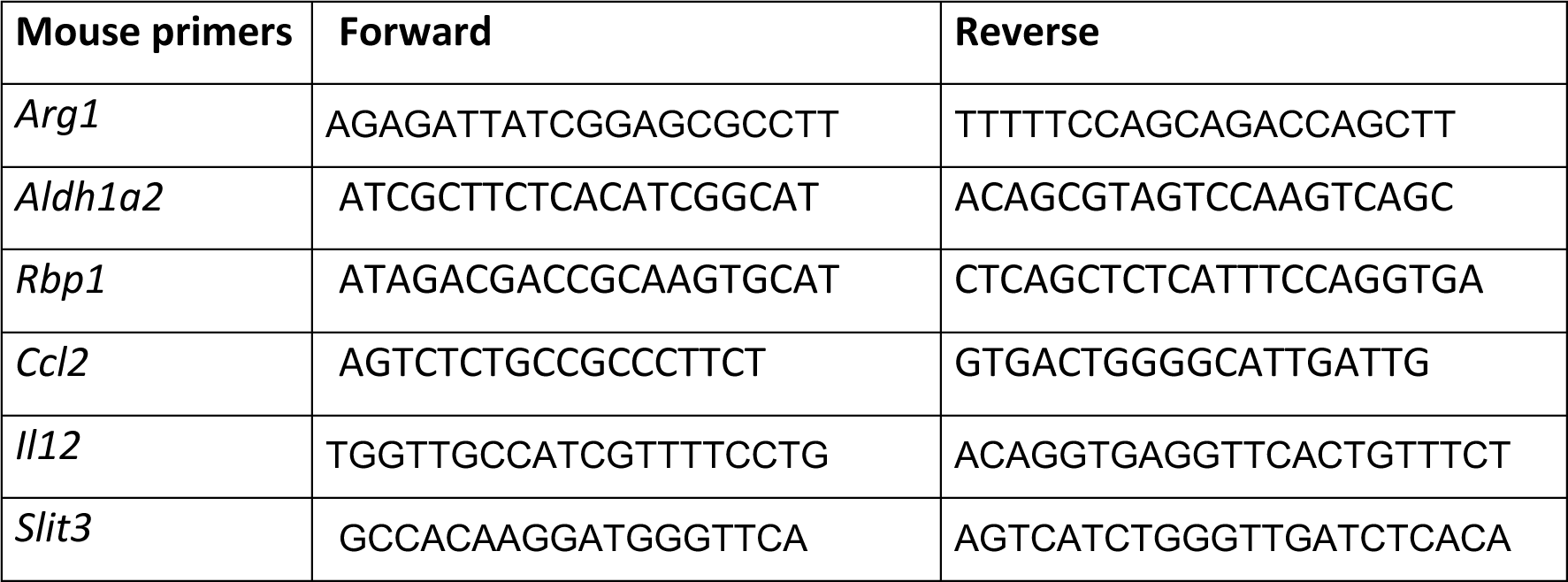
Mouse RT-PCR primers.

## Author contributions

MF designed the whole study and prepared the manuscript with help from all coauthors. MVC and JM designed, performed, and analyzed all nitrofen experiments. RC and RMC designed, performed and analyzed all the Wt1-conditional mice model experiments. JM, LC, BS and MAL purified immunomodulators and tested them in vitro and helped in nitrofen models. LM and JAT contributed to the clinical analysis of the nitrofen-administered animals. All authors contributed with suggestions and critically read the manuscript.

## Acknowledgements.

This research was funded by grants from “Ministerio de Ciencia e Innovación” (SAF2016-75988-R and PID-2019104760RB-100) and “Comunidad de Madrid” (S2017/BMD-3671. INFLAMUNE-CM; FEDER) to MF. The study also received institutional grants from “Fundación Ramón Areces” and “Banco de Santander.”

## Notes

### Competing Interest Statement

The authors have declared no competing interest.

## References

1. Leeuwen L, Fitzgerald DA. Congenital diaphragmatic hernia. Journal of paediatrics and child health 2014; 50(9): 667–73.

2. Chatterjee D, Ing RJ, Gien J. Update on Congenital Diaphragmatic Hernia. Anesth Analg 2020; 131(3): 808–21.

3. Tovar JA. Congenital diaphragmatic hernia. Orphanet J Rare Dis 2012; 7: 1.

4. Greer JJ. Current concepts on the pathogenesis and etiology of congenital diaphragmatic hernia. Respir Physiol Neurobiol 2013; 189(2): 232–40.

5. Kirby E, Keijzer R. Congenital diaphragmatic hernia: current management strategies from antenatal diagnosis to long-term follow-up. Pediatr Surg Int 2020; 36(4): 415–29.

6. Deprest JA, Nicolaides K, Gratacos E. Fetal surgery for congenital diaphragmatic hernia is back from never gone. Fetal Diagn Ther 2011; 29(1): 6–17.

7. Coste K, Beurskens LW, Blanc P, et al. Metabolic disturbances of the vitamin A pathway in human diaphragmatic hernia. Am J Physiol Lung Cell Mol Physiol 2015; 308(2): L147–57.

8. Greer JJ, Babiuk RP, Thebaud B. Etiology of congenital diaphragmatic hernia: the retinoid hypothesis. Pediatric research 2003; 53(5): 726–30.

9. Schreiner Y, Schaible T, Rafat N. Genetics of diaphragmatic hernia. Eur J Hum Genet 2021; 29(12): 1729–33.

10. Yu L, Hernan RR, Wynn J, Chung WK. The influence of genetics in congenital diaphragmatic hernia. Seminars in perinatology 2020; 44(1): 151169.

11. Pasutto F, Sticht H, Hammersen G, et al. Mutations in STRA6 cause a broad spectrum of malformations including anophthalmia, congenital heart defects, diaphragmatic hernia, alveolar capillary dysplasia, lung hypoplasia, and mental retardation. Am J Hum Genet 2007; 80(3): 550–60.

12. Beecroft SJ, Ayala M, McGillivray G, et al. Biallelic hypomorphic variants in ALDH1A2 cause a novel lethal human multiple congenital anomaly syndrome encompassing diaphragmatic, pulmonary, and cardiovascular defects. Hum Mutat 2021; 42(5): 506–19.

13. Dalmer TRA, Clugston RD. Gene ontology enrichment analysis of congenital diaphragmatic hernia-associated genes. Pediatric research 2019; 85(1): 13–9.

14. Nakamura H, Doi T, Puri P, Friedmacher F. Transgenic animal models of congenital diaphragmatic hernia: a comprehensive overview of candidate genes and signaling pathways. Pediatr Surg Int 2020; 36(9): 991–7.

15. Clugston RD, Klattig J, Englert C, et al. Teratogen-induced, dietary and genetic models of congenital diaphragmatic hernia share a common mechanism of pathogenesis. The American journal of pathology 2006; 169(5): 1541–9.

16. Kling DE, Schnitzer JJ. Vitamin A deficiency (VAD), teratogenic, and surgical models of congenital diaphragmatic hernia (CDH). Am J Med Genet C Semin Med Genet 2007; 145C(2): 139–57.

17. Noble BR, Babiuk RP, Clugston RD, et al. Mechanisms of action of the congenital diaphragmatic hernia-inducing teratogen nitrofen. Am J Physiol Lung Cell Mol Physiol 2007; 293(4): L1079–87.

18. Nakazawa N, Takayasu H, Montedonico S, Puri P. Altered regulation of retinoic acid synthesis in nitrofen-induced hypoplastic lung. Pediatr Surg Int 2007; 23(5): 391–6.

19. Jones CV, Ricardo SD. Macrophages and CSF-1: implications for development and beyond. Organogenesis 2013; 9(4): 249–60.

20. Mantovani A, Biswas SK, Galdiero MR, Sica A, Locati M. Macrophage plasticity and polarization in tissue repair and remodelling. The Journal of pathology 2012.

21. Pollard JW. Trophic macrophages in development and disease. Nat Rev Immunol 2009; 9(4): 259–70.

22. Cecchini MG, Dominguez MG, Mocci S, et al. Role of colony stimulating factor-1 in the establishment and regulation of tissue macrophages during postnatal development of the mouse. Development 1994; 120(6): 1357–72.

23. Abood EA, Jones MM. Macrophages in developing mammalian skeletal muscle: evidence for muscle fibre death as a normal developmental event. Acta anatomica 1991; 140(3): 201–12.

24. Correale J, Farez MF. Parasite infections in multiple sclerosis modulate immune responses through a retinoic acid-dependent pathway. Journal of immunology (Baltimore, Md: 1950) 2013; 191(7): 3827–37.

25. Manicassamy S, Ravindran R, Deng J, et al. Toll-like receptor 2-dependent induction of vitamin A-metabolizing enzymes in dendritic cells promotes T regulatory responses and inhibits autoimmunity. Nat Med 2009; 15(4): 401–9.

26. Perveen S, Ayasolla K, Zagloul N, et al. MIF inhibition enhances pulmonary angiogenesis and lung development in congenital diaphragmatic hernia. Pediatric research 2019; 85(5): 711–8.

27. Francisco S, Billod JM, Merino J, et al. Induction of TLR4/TLR2 Interaction and Heterodimer Formation by Low Endotoxic Atypical LPS. Front Immunol 2021; 12: 748303.

28. Carmona R, Canete A, Cano E, Ariza L, Rojas A, Munoz-Chapuli R. Conditional deletion of WT1 in the septum transversum mesenchyme causes congenital diaphragmatic hernia in mice. eLife 2016; 5.

29. Wang YN, Tang Y, He Z, et al. Slit3 secreted from M2-like macrophages increases sympathetic activity and thermogenesis in adipose tissue. Nat Metab 2021; 3(11): 1536–51.

30. Kawai T, Akira S. The role of pattern-recognition receptors in innate immunity: update on Toll-like receptors. Nat Immunol 2010; 11(5): 373–84.

31. Nucera S, Biziato D, De Palma M. The interplay between macrophages and angiogenesis in development, tissue injury and regeneration. Int J Dev Biol 2011; 55(4-5): 495–503.

32. Lang RA, Bishop JM. Macrophages are required for cell death and tissue remodeling in the developing mouse eye. Cell 1993; 74(3): 453–62.

33. Klimchenko O, Di Stefano A, Geoerger B, et al. Monocytic cells derived from human embryonic stem cells and fetal liver share common differentiation pathways and homeostatic functions. Blood 2011; 117(11): 3065–75.

34. Rae F, Woods K, Sasmono T, et al. Characterisation and trophic functions of murine embryonic macrophages based upon the use of a Csf1r-EGFP transgene reporter. Dev Biol 2007; 308(1): 232–46.

35. Martin P, D’Souza D, Martin J, et al. Wound healing in the PU.1 null mouse--tissue repair is not dependent on inflammatory cells. Curr Biol 2003; 13(13): 1122–8.

36. Jones CV, Williams TM, Walker KA, et al. M2 macrophage polarisation is associated with alveolar formation during postnatal lung development. Respir Res 2013; 14: 41.

37. Seledtsov VI, Seledtsova GV. A balance between tissue-destructive and tissue-protective immunities: a role of toll-like receptors in regulation of adaptive immunity. Immunobiology 2012; 217(4): 430–5.

38. Manicassamy S, Pulendran B. Modulation of adaptive immunity with Toll-like receptors. Semin Immunol 2009; 21(4): 185–93.

39. Jalian HR, Liu PT, Kanchanapoomi M, Phan JN, Legaspi AJ, Kim J. All-trans retinoic acid shifts Propionibacterium acnes-induced matrix degradation expression profile toward matrix preservation in human monocytes. J Invest Dermatol 2008; 128(12): 2777–82.

40. Pulendran B, Tang H, Manicassamy S. Programming dendritic cells to induce T(H)2 and tolerogenic responses. Nat Immunol 2010; 11(8): 647–55.

41. Holder AM, Klaassens M, Tibboel D, de Klein A, Lee B, Scott DA. Genetic factors in congenital diaphragmatic hernia. Am J Hum Genet 2007; 80(5): 825–45.

42. Gilbert RM, Gleghorn JP. Connecting clinical, environmental, and genetic factors point to an essential role for vitamin A signaling in the pathogenesis of congenital diaphragmatic hernia. Am J Physiol Lung Cell Mol Physiol 2023; 324(4): L456–L67.

43. Mendelsohn C, Lohnes D, Decimo D, et al. Function of the retinoic acid receptors (RARs) during development (II). Multiple abnormalities at various stages of organogenesis in RAR double mutants. *Development (Cambridge*, England) 1994; 120(10): 2749–71.

44. Quadro L, Blaner WS, Salchow DJ, et al. Impaired retinal function and vitamin A availability in mice lacking retinol-binding protein. EMBO J 1999; 18(17): 4633–44.

45. Beurskens LW, Tibboel D, Lindemans J, et al. Retinol status of newborn infants is associated with congenital diaphragmatic hernia. Pediatrics 2010; 126(4): 712–20.

46. Clugston RD, Zhang W, Alvarez S, de Lera AR, Greer JJ. Understanding abnormal retinoid signaling as a causative mechanism in congenital diaphragmatic hernia. Am J Respir Cell Mol Biol 2010; 42(3): 276–85.

47. Balounova J, Vavrochova T, Benesova M, Ballek O, Kolar M, Filipp D. Toll-like receptors expressed on embryonic macrophages couple inflammatory signals to iron metabolism during early ontogenesis. Eur J Immunol 2014; 44(5): 1491–502.

48. Toelen J, Carlon M, Claus F, et al. The fetal respiratory system as target for antenatal therapy. Facts Views Vis Obgyn 2011; 3(1): 22–35.

49. Eiro N, Pidal I, Fernandez-Garcia B, et al. Impact of CD68/(CD3+CD20) ratio at the invasive front of primary tumors on distant metastasis development in breast cancer. PloS one 2012; 7(12): e52796.

50. Pries AR, Reglin B, Secomb TW. Remodeling of blood vessels: responses of diameter and wall thickness to hemodynamic and metabolic stimuli. Hypertension 2005; 46(4): 725–31.

51. Guerrero NA, Camacho M, Vila L, et al. Cyclooxygenase-2 and Prostaglandin E2 Signaling through Prostaglandin Receptor EP-2 Favor the Development of Myocarditis during Acute Trypanosoma cruzi Infection. PLoS Negl Trop Dis 2015; 9(8): e0004025.

52. Cuervo H, Guerrero NA, Carbajosa S, et al. Myeloid-Derived Suppressor Cells Infiltrate the Heart in Acute Trypanosoma cruzi Infection. J Immunol 2011; 187(5): 2656–65.

