## Supplemental figures for "Anti-inflammatory immunomodulation for the treatment of Congenital Diaphragmatic Hernia"

Supplemental data

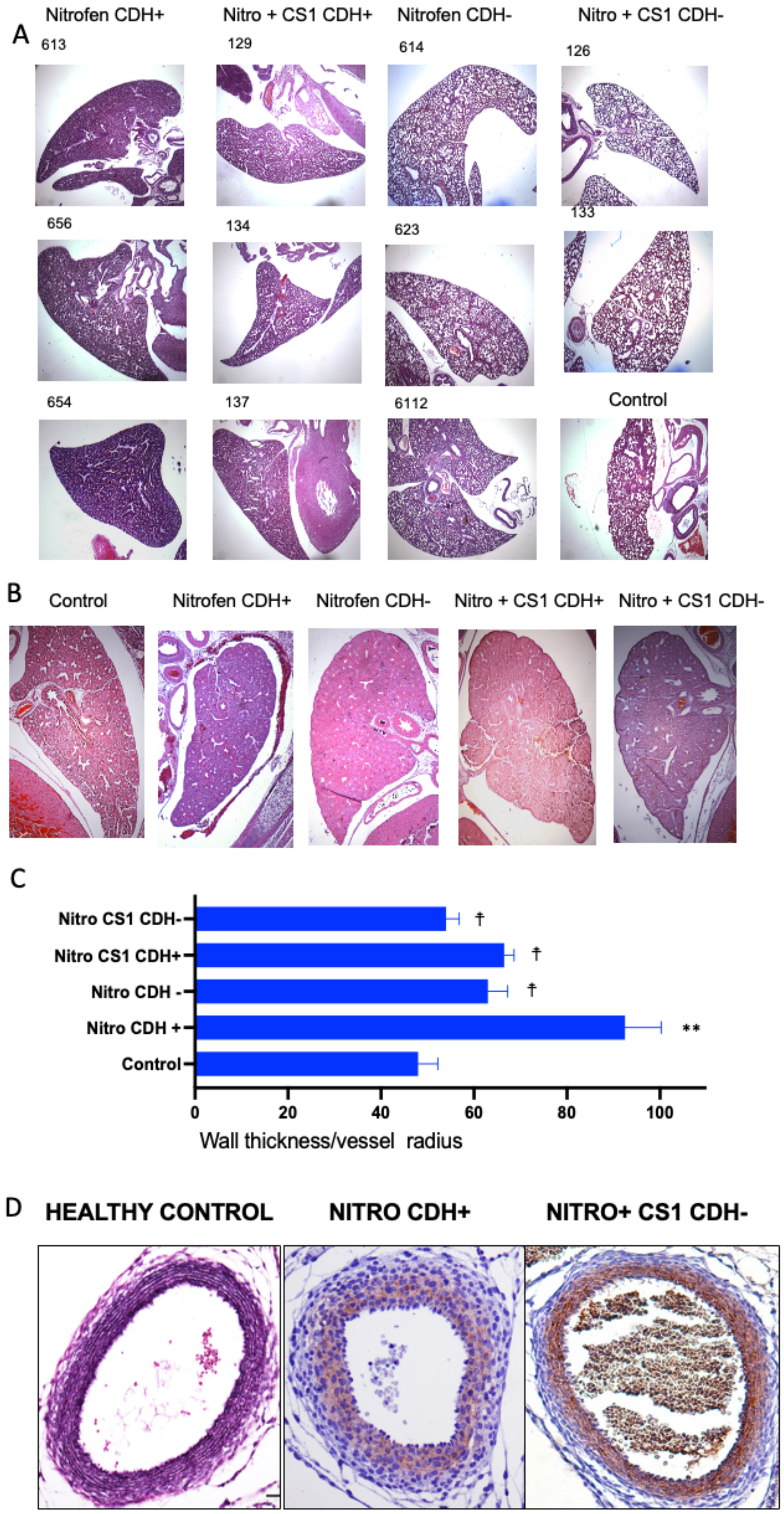

**Figure S1. Effect of CS1 ligand in alteration in lung development induced by nitrofen.** Pregnant Wistar rat females were administered nitrofen , or olive oil (control), at E9.5 and 3 days later injected intraperitoneally with 10 mg/Kg CS1. A) Hematoxylin-eosin images of E21 lungs of 3 different fetuses from the different groups. B) A representative image of each group at E18. C) Wall thickness/vessel radius ratio. Analysis of E18 lung vessels from the different groups. 3 independent calibrations for different lung sections of 3 different foetuses were staining with van Giemson; ** = p>0.001, respect to control normal group; † = p>0.001 respect to untreated CDH+ nitrofen group. D) Representative van Giemson staining of the indicated groups.

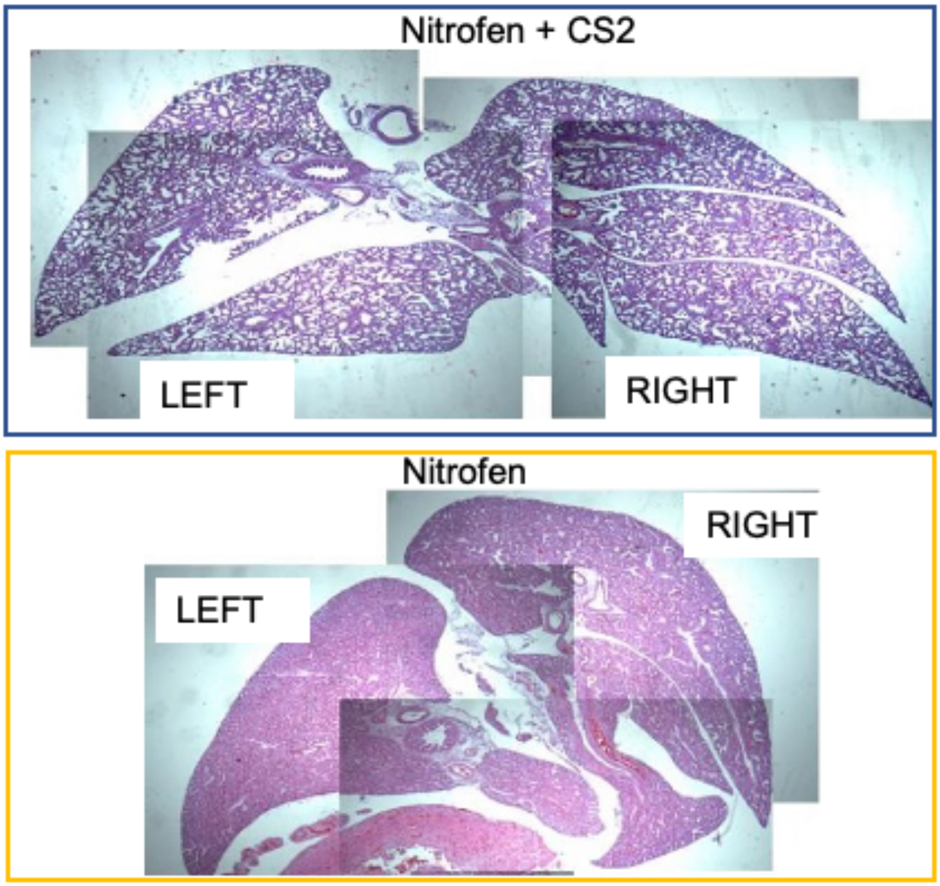


**Figure S2 Effect of CS2 ligand in alteration in lung development induced by nitrofen**. Pregnant Wistar rat females were administered nitrofen at E9.5 and 3 days later injected intraperitoneally with 100 mg/Kg CS2 Superposed images to reconstruct the complete lungs from Nitrofen or Nitrofen (CDH+) and CS1 treated (CDH-) animals at E21.

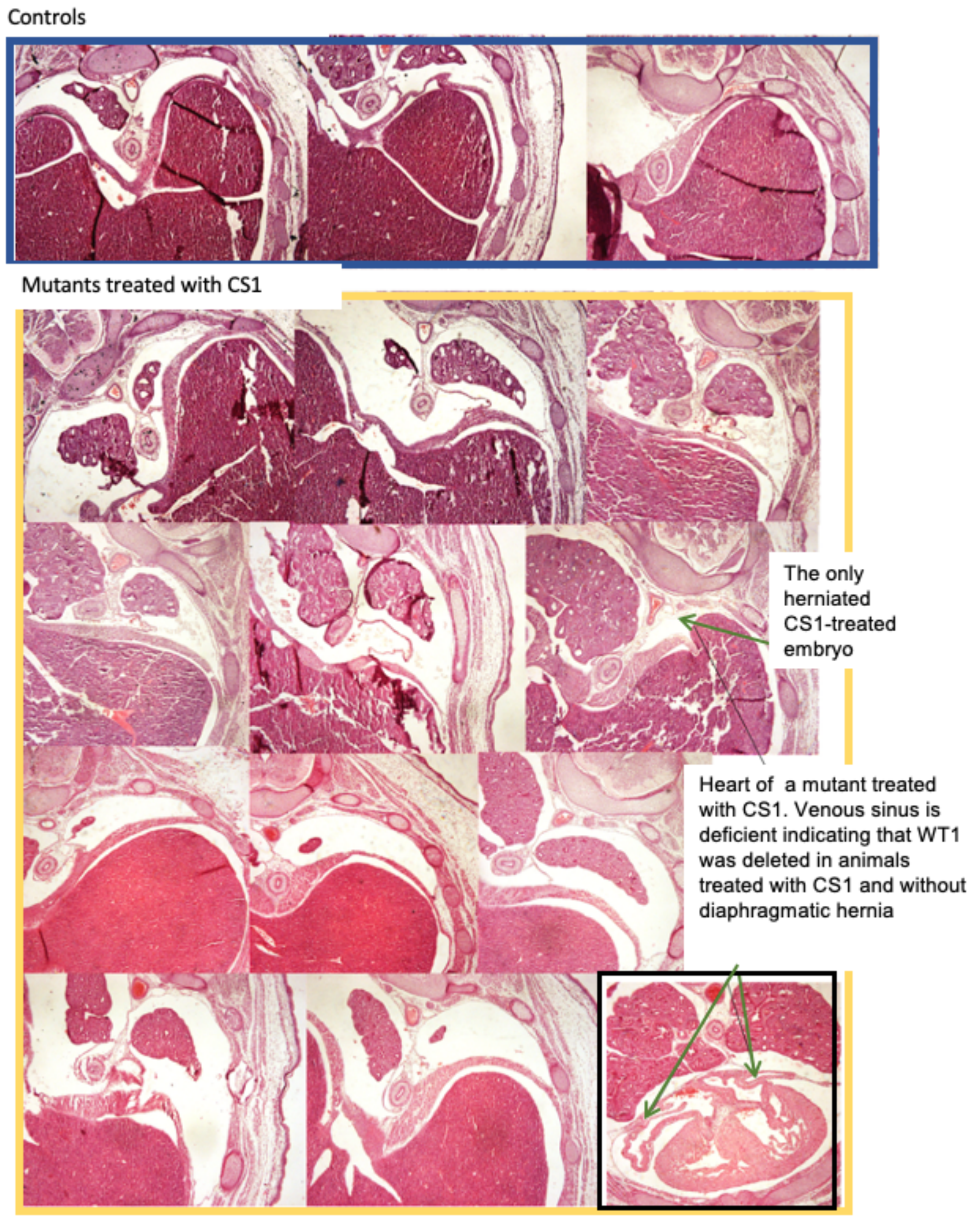


**Figure S3 Effect of CS1 ligand in diaphragmatic hernia of G2-GATA4$^{Cre}$;Wt1$^{fl/fl}$ mice .** Pregnant mice mothers were treated with CS1 or PBS intraperitoneally twice at E9.5 and E10.5. Embryos were analysed at E15.5. Images of the diaphragm from the 11 WT1 mutant embryos of mothers treated CS1 obtained. The heart images of a mutant CDH- is also shown. Embryos (3) from control mothers are also shown.

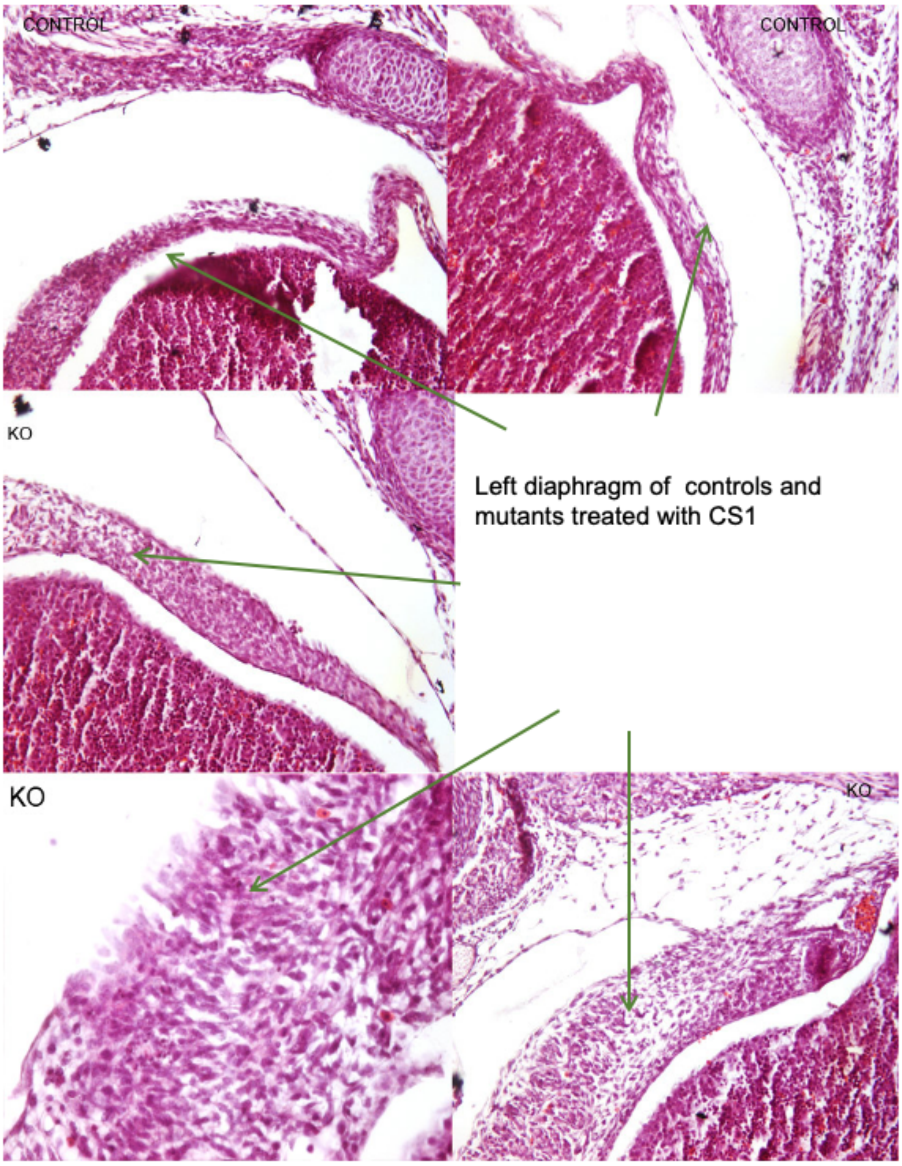


**Figure S4 Effect of CS1 ligand in diaphragmatic hernia of G2-GATA4$^{Cre}$;Wt1$^{fl/fl}$ mice**. Pregnant mice mothers were treated with CS1 or PBS intraperitoneally twice at E9.5 and E10.5. Embryos were analysed at E15.5. Representative images of the diaphragm from Wt1-mutant embryos of mothers treated with CS1 (3) and from Wt1-wild type control embryos from CS1-treated mothers (2) are shown.
